# Shift work dynamics and division of labor: honeybee foraging and fanning tasks

**DOI:** 10.1101/2020.08.17.254755

**Authors:** Manuel A. Giannoni-Guzmán, Tugrul Giray, Jose L. Agosto-Rivera

## Abstract

In complex societies common social needs such as vigilance, care giving, resource gathering, and production are attended around the clock. In humans, these services are constantly provided using a shift work strategy where different individuals, or groups of individuals perform their tasks at different times of the day. However, shift work strategy in job organization in other social organisms remains unclear. Previous studies in honeybees for two jobs support shift work for only pollen foragers and not for nursing behavior. Here we examined shift work dynamics for three types of jobs performed by honeybee foragers. Specifically, we studied pollen foragers, non-pollen foragers and bees fanning at the entrance of the colony, a job important for orientation and temperature control. Major features of the observed shift work were: 1) individuals can be divided into early and late shifts; 2) there are constant workers; 3) based on job, shift work is performed by fewer or greater number of individuals; 4) shift work of an individual is plastic and may change with age; 5) foraging and fanning shifts are coupled yet dissociable. This study adds to the findings that shift work is not exclusive to modern human societies and that a natural form of shift work exists in honeybees. These results suggest that shift work in honeybees is a feature of worker division of labor. Future studies aiming to further understand the structure, function and mechanism of this natural form of shift work in honey bees not only could have an impact on agriculture but also may provide insight into alternative forms of shift work strategies that may reduce the various health problems associated with shift work in humans.

## Introduction

Principles that organize social work are common across social organisms (Gordon, 2007; Oster & Wilson, 1978). Specialization, based on ontogenetic, morphological or genetic mechanisms, occurs in many social species (Fjerdingstad & Crozier, 2006; Jeanson, Fewell, Gorelick, & Bertram, 2007; O’Riain, Jarvis, Alexander, Buffenstein, & Peeters, 2000; Robinson, 1992). Parallel processes performed by multiple agents result in networks. Networks of individuals can modulate behavior via feedback regulation, that may depend on order of task performance, such as foraging followed by nectar unloading and storage (Craig et al., 2012; Jeanne, 1986; O’Donnell & Jeanne, 1992) or based on chemical communication such as pheromones or cuticular hydrocarbons (Inoshita, Martin, Marion-Poll, & Ferveur, 2011; Pankiw, 2004; Sagili, Pankiw, & Metz, 2011). Spatial organization confines activities to specific locations, often enhancing the effects of other mechanisms that organize work (Jandt & Dornhaus, 2009; Mersch, Crespi, & Keller, 2013). Temporal organization, restricts the performance of a task to a specific time period of the day and may have molecular, cellular and behavioral correlates (C S Pittendrigh, 1993; Roenneberg, Wirz-Justice, & Merrow, 2003; Southerton, 2006). At the behavioral level, various temporal organization strategies have emerged throughout history. Among them, shift work strategies have become a mainstay in modern human societies (Folkard, 2003a; Pati, Chandrawanshi, & Reinberg, 2001). However, shift work has not been studied extensively in social insects.

Shift work is a method of organizing individuals or groups to perform specific tasks at different times of the day such that these tasks can be continuously performed (IARC Working Group on the Evaluation of Carcinogenic Risks to Humans, 2010; Pati et al., 2001). Professions such as health care, emergency response teams (e.g. firemen), transportation and food service use various shift work strategies to provide these essential services around the clock (Adan et al., 2012). Although shift work strategies succeed by providing many benefits for society and employers, there are costs at the individual and social level. Catastrophes such as the Chernobyl nuclear meltdown, Three Mile Island and the Exxon Valdez oil spill, have been linked to errors associated with shift work (Akerstedt & Wright, 2009; Folkard, 2003b; Klerman, 2005; Mitler et al., 1988; Pati et al., 2001; S. M W Rajaratnam & Arendt, 2001). Studies examining the relationship of shift work and health problems provide convincing evidence that misalignment of circadian rhythms is one of the key contributors to many, if not all, of the negative effects associated with shift work (Knutsson, 1989; Shantha M W Rajaratnam, Howard, & Grunstein, 2013). It has thus become important to study what strategies other social organisms, use to achieve their 24/7 needs.

In honeybees, colony structure is best defined by castes with clear division of labor system with diploid queens and haploid males (drones) attending reproductive tasks, while sterile diploid workers perform all other jobs associated with colony maintenance (Robinson, 1992; Mark L Winston, 1987). Within workers, division of labor is an age-related process, where workers perform a series of tasks from the moment they emerge as adults and change tasks as they age until they begin foraging (∼21 days of age) (Seeley, 1985, 1995; Mark L Winston, 1987). The rate of division of labor in workers has been shown to be genetically, behaviorally and hormonally regulated and as a result, individuals of the same age can be observed performing different tasks (Giray & Robinson, 1996; Giray, Guzman-Novoa, et al., 1999; Huang & Robinson, 1992; Leoncini et al., 2004).

In the colony tasks such as brood care, cleaning cells, fanning and foraging are performed throughout the day, or for extended periods of time. Whether individuals are constantly performing these tasks or if they use shift work strategies has been asked. Previous work examined if nurses used shift work or similar strategies to take care of the brood. Moore and colleagues (1998) marked and observed task performance of individual bees and found that brood care is performed throughout the day without specific timing (Moore, Angel, Cheeseman, Fahrbach, & Robinson, 1998). This coincides with the idea that the brood releases pheromones that make honey bee workers lose their circadian rhythmicity such that they feed the brood (Moore, 2001; Nijland & Hepburn, 1985; Yair Shemesh, Eban-Rothschild, Cohen, & Bloch, 2010; Spangler, 1972; Stussi, 1972). Based on these studies nurses take care and feed the brood, in a similar manner as human mothers take care of new-born children, around the clock. However, this finding in brood care may not extend to other jobs in the honeybee colony.

Fanning behavior is a task that workers perform to regulate the temperature of the colony, release Nasonov’s pheromone and mature honey (Seeley, 1995; Mark L Winston, 1987). A study examining thermoregulation of the colony, focusing of fanning behavior, found that colonies with a uniform genetic background (originated from one male) are less efficient at maintaining temperature levels compared to colonies with a diverse genetic background (J C Jones, Myerscough, Graham, & Oldroyd, 2004). However, whether bees use a particular strategy to organize fanning throughout the day has remained unexplored.

In the case of foraging, bees use the full daylight period in order to gather the various resources that colonies need on a daily basis. Through the use of sun compass navigation (R M Goodwin & Lewis, 1987; von Frisch, 1967), time memory (Moore & Doherty, 2009; Moore, Van Nest, & Seier, 2011; B. N. Van Nest & Moore, 2012) and circadian rhythms(Bloch & Robinson, 2001; Cheeseman et al., 2012; Eban-Rothschild & Bloch, 2012; Y Shemesh, Cohen, & Bloch, 2007; Yair Shemesh et al., 2010), bees predict the availability of different resources throughout the day. Individual workers can specialize in the collection of a specific resource such as pollen, nectar or water (Fewell & Page, 1993; Robinson & Page, 1989; Seeley, 1995). Studies examining the underlying factors of this resource specialization have found genetic, neuroendocrine and behavioral differences between pollen and nectar specialists (Barron, Maleszka, Vander Meer, & Robinson, 2007; Erber, Hoormann, & Scheiner, 2006; Giray, Galindo-Cardona, & Oskay, 2007; Page & Erber, 2002; Scheiner, Barnert, & Erber, 2003; Scheiner, Page, & Erber, 2001; Scheiner, Plückhahn, Oney, Blenau, & Erber, 2002; Scheiner, Toteva, Reim, SÃ,vik, & Barron, 2014; Taylor, Robinson, Logan, Laverty, & Mercer, 1992; Wagener-Hulme, Kuehn, Schulz, & Robinson, 1999).

In contrast to brood care, in a recent study, researchers captured incoming pollen foragers in the morning and afternoon for four days and genotyped them with microsatellite DNA markers (Kraus, Gerecke, & Moritz, 2011). They uncovered that a small percentage of pollen foragers from specific patrilines were only captured in the morning, while foragers from other patrilines were only captured in the afternoon. This finding suggests that some pollen foragers make their foraging trips in shifts and this behavior is in part influenced by the genetic origin of the individual (Kraus et al., 2011). Organization of shift work for pollen foraging and potentially other jobs can be examined through direct observations as was done for nursing.

Here we present a comprehensive analysis of foraging and fanning behavior in honeybee workers to determine the presence and organization of shift work. In this study, we determined whether a shift work strategy is evident in pollen, non-pollen foraging and behaviors and if so, 2) describe the behavioral characteristics of this shift work. We performed direct behavioral observations at the hive entrance workers of an age cohort. Our central hypothesis was that if foragers perform specific tasks in shifts then we would observe groups of individuals performing these behaviors at specific times of the day. To address specific characteristics of shift work we conducted our observations over most of the foraging life of the age cohort. In this way, we could examine the degree of plasticity associated with shift work and whether the organization of shift workers varies between different jobs. Lastly, we examined possible relationships of the temporal allocation between foraging and fanning tasks for each individual.

## Materials and Methods

### 1. Observation ramp

A two-story hive with a naturally mated queen was fitted with an extended entrance ramp with a glass top measuring 45cm wide and 40cm in length (Giray et al., 2007). Briefly, to train the bees to the entrance ramp, we first installed the ramp without the glass top. Two days following the placement of the ramp a piece of glass of 5cm in length was lined with colored tape and placed in the ramp. The following days the length of the glass was slowly extended until it covered the full length of the ramp. The glass top assured a narrow space within which bees were unable to cover each other or walk upside down.

### 2. Bees

Honeybee workers were obtained from 2 healthy colonies (collection colonies) with a naturally mated queen at the University of Puerto Rico Bee Research Facility at the Gurabo Experimental Agriculture Station. From each of the colonies, we marked three groups of 500 bees (n=3,000 marked individuals) with a three-day interval between each marking group. To mark, we extracted 2 brood frames with large numbers of capped brood in the afternoon. The frames were gently brushed to remove the attending nurses and transported to our laboratory incubator (Thermo Scientific Precision Incubator 815), where they remained overnight. Bees that had emerged on the following morning were extracted and individualized by applying a colored numbered tag in the thorax and a paint dot (acrylic, Testors^®^: TES1127TT, TES1146TT and TES1172TT) in the abdomen identifying the Age cohort. After marking, bees were placed inside of the colony that had been previously fitted with the observation ramp.

### 3. Observation periods

Observations were twice a day for 14 days, from 9:00-11:00 and from 14:00-16:00, in a similar manner as in a previous study (Krauss et al., 2011). In addition, these periods were chosen to prevent the overlap of foraging trips between observation periods. Researchers have observed that the duration of foraging trips can range from 4-25 min on average but foraging trips longer than 50 minutes have been recorded ((Mattu, Raj, & Thakur, 2012; Partap, Shukla, & Verma, 2000; Singh, 2009; Wagner, Van Nest, Hobbs, & Moore, 2013). Before each observation period began, a thin coating of petroleum jelly (Vaseline^®^) was applied to the glass to prevent bees from walking upside down. Colonies were observed sequentially during the summer, in this manner colony 1 observations took place from May 25^th^ – June 7^th^, 2012 while colony 2 observations took place from June 28^th^ – July 11^th^, 2012. During observation the entry, exit and fanning behavior of each individual was recorded in a laboratory notebook with an accompanying time stamp, and later transcribed to JMP for data analysis.

During the 14-day observation periods, for colony 1, of the 1,500 marked individuals we were able to observe a total of 1,030 bees and recorded 5,102 individual observations. For the same duration, 535 of 1,500 marked bees were observed in colony 2 and a total of 2,698 individual observations were recorded. Observations for colony 2 took place during Puerto Rico’s rainy season, and constant interruptions due to weather conditions may account for the differences in the number of observations. Our methodology allowed us to record, on average5 direct behavioral observation for each of more than a thousand individuals.

### 4. Morningness ratio

To establish if bees perform shift work for each of the observed behaviors (foraging trips or fanning) we tabulated the number of observations during the morning observation periods and afternoon observation periods for each individual. We then calculated the ratio of morning observations over the total observations. This formula was modified from that previously described and used by Moore et al. (1998). In this manner, individuals that mainly forage or fan in the afternoon would have morningness ratios close to 0 (afternoon shift), while those that forage or fan mainly in the morning would have a ratio close to 1 (morning shift). Similarly, if individuals have no temporal preference for performing a specific task, they would have a ratio close to 0.5 (no shift).

### 5. Foraging patterns

To answer if bees’ preference to forage in the morning or afternoon changed as they aged, we examined each individual’s foraging trip observations in scatterplots. We identified five foraging pattern phenotypes: morning; afternoon; morning-afternoon, who began in the morning and after some time switched to the afternoon; afternoon-morning, began in the afternoon and switched in the morning; and constant. For an individual to be included in a foraging pattern their observations had to span for a period of 6 days or more and the majority of these had to have occurred within the 12-19 days of age to control for any possible bias fewer observations on an individual may generate.

### 6. Data Analysis

For both foraging trips and fanning behavior only individuals with 3 or more observations were considered for data analysis. We also excluded individuals for whom all observations were taken on the same day. Comparison of the observed frequency distributions of the morningness ratio for foraging trips and fanning behavior, for each colony, was statistically compared using chi-square goodness of fit with theoretical frequencies from a binomial distribution that assumes no shift work (null hypothesis). To compare the observed distributions of each colony (foraging trips or fanning) we utilized the Kolmogorov-Smirnov test of distributions. Median test was used to compare the foraging pattern frequency distributions, the mean trips taken, the probability of taking a foraging trip and the mean number of trips in a foraging period. For the correlations of the foraging and fanning morningness ratios, pairs of observations from the same day were tested with Kendall’s tau association test. All statistical analyses were performed using the statistical software program JMP (SAS Institute Inc.). Figures were prepared using GraphPad PRISM 6.00, GraphPad software, La Jolla California USA and R (R Core Team).

## Results

### Foragers use two temporal strategies to gather resources for the hive

To determine if all foragers go out throughout the day or if groups of individual bees forage at different times of the day, we calculated the number of morning observations over the total number of observations (morningness ratio) for each forager. Since foraging in African-hybrids, such as the ones used in this study, can start as early as 11 days of age (Giray, Huang, Guzman-Novoa, & Robinson, 1999; M L Winston, 2003; Mark L Winston, 1987), we used the data observations from 17 days of age onward. Consistent with our hypothesis, our results revealed that more than 40% of the individuals exclusively foraged either in the morning or afternoon, now on referred to as shift workers (Figure 1A). In addition, to shift workers, we also observed constant workers, which foraged both in the morning and afternoon. To determine if the observed shift work ratios were significantly different from chance, the observed distribution was compared with a theoretical binomial distribution that assumed the absence of shifts. This comparison using Pearson’s *X*^*2*^ resulted in significant differences for both colonies sampled, suggesting that groups of workers forage at different times of the day (colony 1: Pearson’s *X*^*2*^*= 1009*.*53, p<< 0*.*01*, n=227; colony 2: Pearson’s *X*^*2*^*= 647*.*73, p << 0*.*01*, n=142). Statistical comparison using the Kolmogorov-Smirnov statistical analysis was also performed to compare the observed distributions of the sampled colonies. This resulted in significant differences between the observed distributions of each colony (Kolmogorov-Smirnov two-sided test, D=0.1671, *p=0*.*02)*, suggesting possible colony-colony differences in shift work.

**Figure 1.**
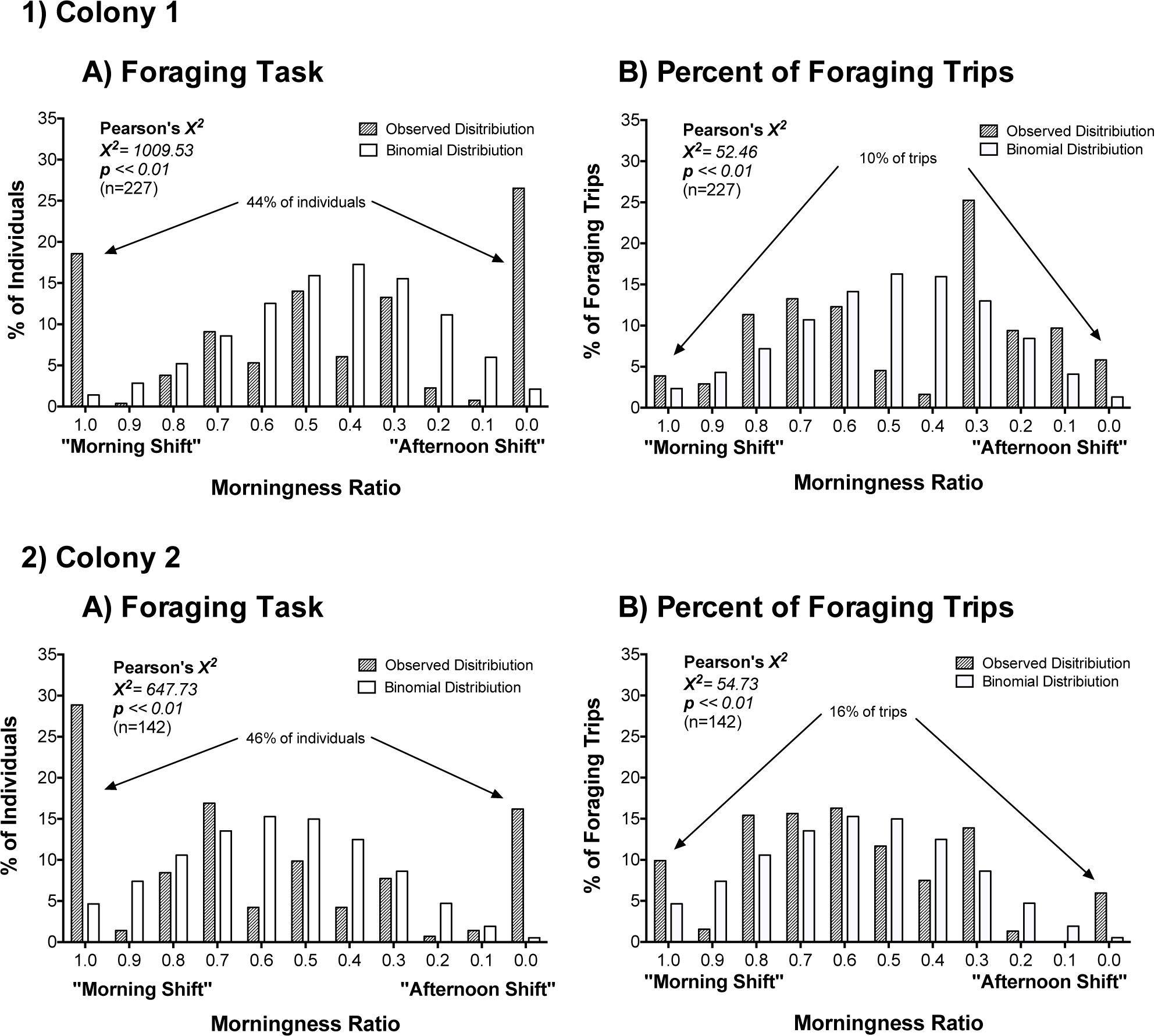
Exclusive shifts in morning and afternoon are present foraging task. **(A)** Frequency distribution of observed morningness ratio for per cent of individuals (shaded bars) compared to a theoretical binomial distribution (white bars) revealed that more than 40% of sampled individuals foraged exclusively in the morning or afternoon as pointed as pointed by arrows for **1)** colony 1 and **2)** colony 2. Goodness of fit test revealed significant differences between the observed and theoretical distributions (colony 1: Pearson’s *X*^*2*^*= 1009*.*53, p<< 0*.*01*, n=227; colony 2: Pearson’s *X*^*2*^*= 647*.*73, p << 0*.*01*, n=142). Comparison between the observed distributions of individuals for each colony via Kolmogorov-Smirnov two-tailed test revealed significant differences between the observed morningness ratio distributions (D=0.17, *p=0*.*02*). **(B)** Frequency distribution of morningness ratio and the present of trips observed (shaded bars) reveals that less than 20% of trips are made by foragers who exclusively forage in the morning or afternoon as pointed. Comparison of each of the observed distribution with a theoretical binomial distribution (white bars) revealed significant differences between the observed and theoretical distributions (colony 1: Pearson’s *X*^*2*^*= 52*.*46, p<< 0*.*01*, n=227; colony 2: Pearson’s *X*^*2*^*= 54*.*73, p << 0*.*01*, n=142). Comparison between the per cent of trips for each colony via the Kolmogorov-Smirnov two-tailed test revealed significant differences each observed distribution (D=0.31, p<<0.01).

Further examination of our data set revealed that the number of observations between shift workers and constant workers varied greatly. We hypothesized that constant workers, who forage throughout the day, would perform at least twice the foraging trips than shift workers, who only forage at specific times of the day. To test this hypothesis, we took into account the number of observation periods, that constant workers would be observed in both periods and the proportion of constant workers that were observed we predicted that constant workers would be responsible for ∼75% of the observed foraging trips. Consistent with our prediction, constant workers account for more than 80% of our observed foraging trips, while shift workers performed less than 20% of foraging trips observed (Figure 1B).

### Shift workers within pollen foragers represent a small subset of individuals

Previous work presenting genetic evidence for shift work in foragers was restricted to pollen foragers (Kraus et al., 2011). In our experiments we observed marked foragers in general and were able to discern between pollen and non-pollen foragers. By separating pollen and non-pollen foragers we found within both pollen and non-pollen foragers there are individuals foraging in shifts (Figure 2). In the case of pollen foragers less than 10% perform foraging in shifts, which is consistent to the genetic work previously published (Kraus et al., 2011). Conversely, 36% percept of non-pollen individuals exclusively forage in the morning or afternoon (Figure 2), suggesting that non-pollen foraging has a stronger shift worker component than pollen foraging.

**Figure 2.**
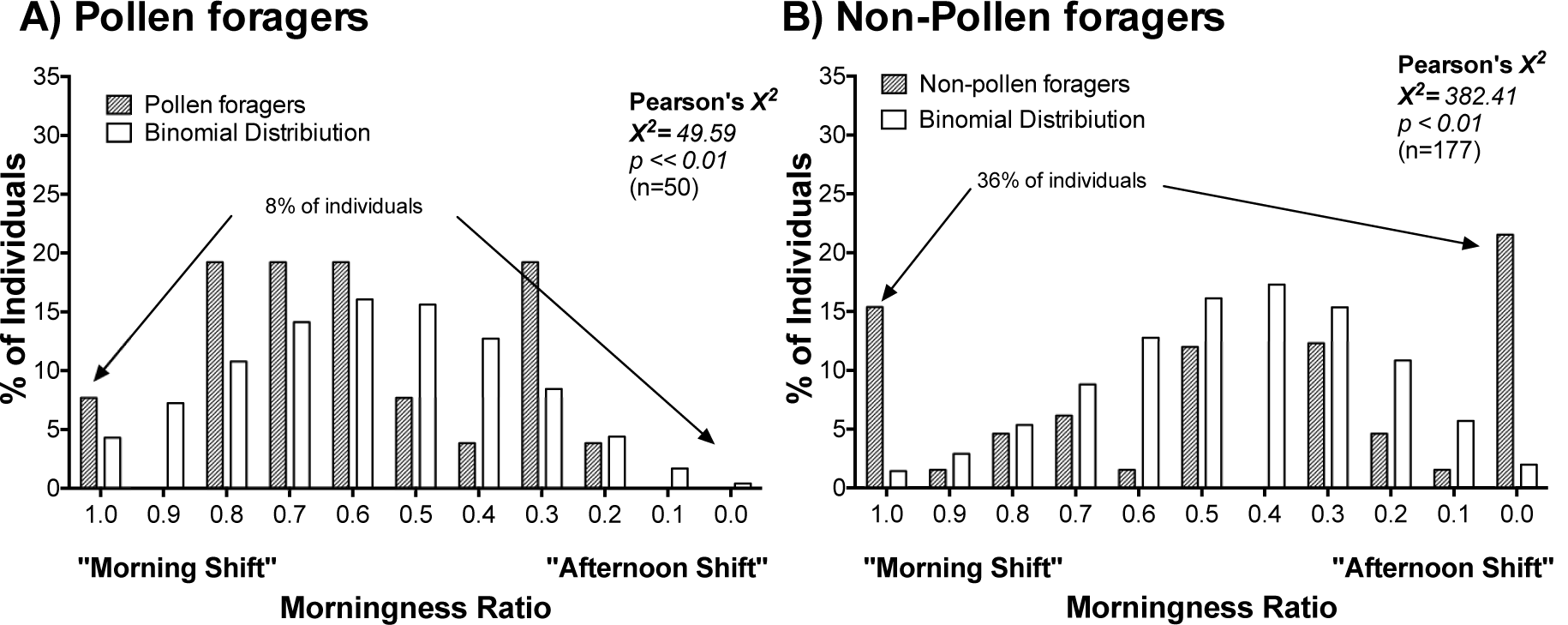
Shift work allocation depends on foraging specialization were only 8% of pollen foragers present a shift. **A)** Frequency distribution of observed morningness ratio for pollen specialists of colony 1 (shaded bars) compared to a theoretical binomial distribution (white bars) of the null hypothesis. Goodness of fit test reveals significant differences between the observed and theoretical distributions (Pearson’s *X*^*2*^*= 49*.*59, p<< 0*.*01*, n=50). **B)** Frequency distribution of observed morningness ratio of non-pollen specialists of colony 1 (shaded bars) compared to a theoretical binomial distribution (white bars) of the null hypothesis (Pearson’s *X*^*2*^*= 382*.*41, p<< 0*.*01*, n=117). Pollen specialists compose less than 20% of individuals that perform foraging exclusively in the morning or afternoon

### Foraging shifts may change as bees age

Since division of labor in honey bee workers is a complex age based process (Seeley, 1985, 1995; Mark L Winston, 1987), we hypothesized that age-related plasticity may be evident in worker shifts. Our approach to address this interest was to examine those individuals for which data was collected over a 6-day time period. Our analysis described five distinct behavioral patterns, which we sorted into different groups: 1) individuals that preferred to forage during one of the periods (morning or afternoon), classified as static shift workers, and 2) individuals that foraged indiscriminately in either period, classified as constant workers (Figure 3 A). In addition, a third foraging pattern was observed, where individuals presented a shift and after some time changed from that shift to the opposite and classified as changing shift workers (morning-afternoon, afternoon-morning) (Figure 3A). Comparing the frequency of each of the foraging patterns shows that constant workers represent more than 60% of the observed foraging population (Figure 3B). We further studied individuals who changed shifts to establish if there was a specific time window in the forager’s life for this change and whether the nature of this change in shift is endogenous or exogenous in origin. By establishing the age at which each of the observed individuals changed shift we were able to establish the age range that presents the highest probability a forager changes shift (Figure 3C). Our results revealed that approximately 75% of individuals change shifts from 11-19 days of age, the early stage of the individuals foraging life.

**Figure 3.**
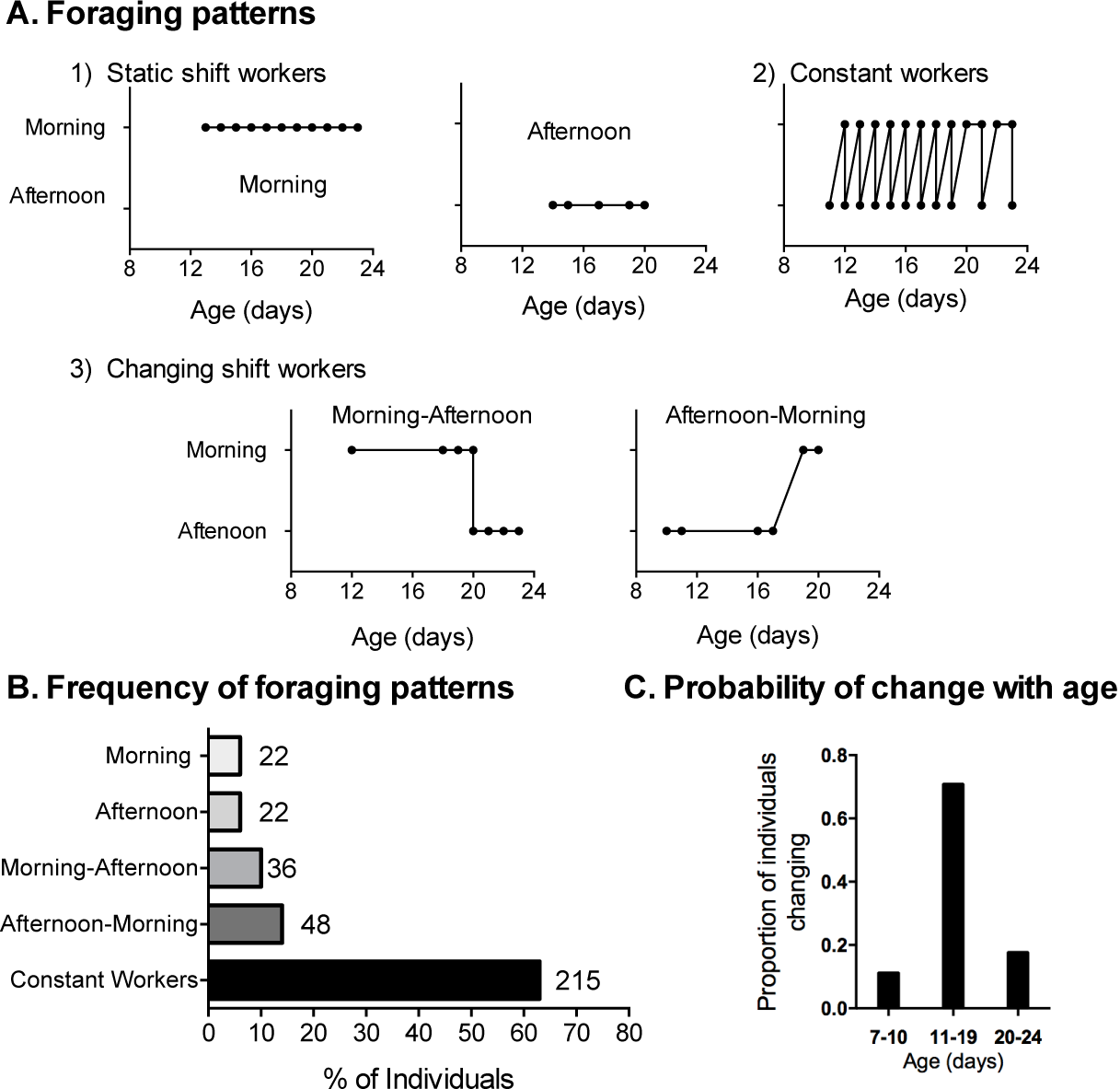
Shift work in foraging is plastic and can change with age. **A)** Examples of the 5 foraging patterns obtained from honeybee entry and exit data from entrance observations: 1) Static Shifts (Morning and Afternoon); 2) Changing shifts (Morning-Afternoon and Afternoon-Morning); and 3) Constant foragers. **B)** Proportion of individuals changing shifts (morning to afternoon or afternoon to morning) at different age blocks. No significant differences were found between the colonies. **C)** Foraging pattern distribution of sampled individuals. Non selective individuals makeup more than 60% of our sample group, while approximately 20-25% of individuals change shifts once during their life and around 15% of individuals have static shifts. Comparison between colonies via contingency analysis did not reveal significant differences.

### Fanning is performed in shifts

While our results show that foraging is performed in shifts in honeybee colonies, whether the observed shift work is endogenously driven or regulated by environmental factors, such as flowers, was not distinguishable in our data set. For this reason, we analyzed if fanning behavior at the entrance of the colony was done in shifts. Given the narrow regulation of temperature in honeybee colonies we hypothesized that fanning behavior at the entrance of the colony would be performed by some individuals in shifts and by others constantly throughout the day. Consistent with this hypothesis, our results show that some workers perform fanning in shifts, while others were observed fanning throughout the day (Figure 4). Comparison between the theoretical binomial distribution for no shift work and the observed distribution via Pearson’s *X*^*2*^ resulted in significant differences for both colonies sampled (colony 1: *X*^*2*^*= 258*.*91, p < 0*.*001, n=45*; colony 2: *X*^*2*^*= 529*.*69 p < 0*.*001, n=22*; Figure 4). In addition, comparison between colonies via Kolmogorov-Smirnov test resulted in significant differences between the observed distributions for the colonies (D = 0.346, p ≤ 0.05). The finding that fanning is also performed in shifts and colonies differ in distribution of individuals, suggest that shift work may be endogenously driven.

**Figure 4.**
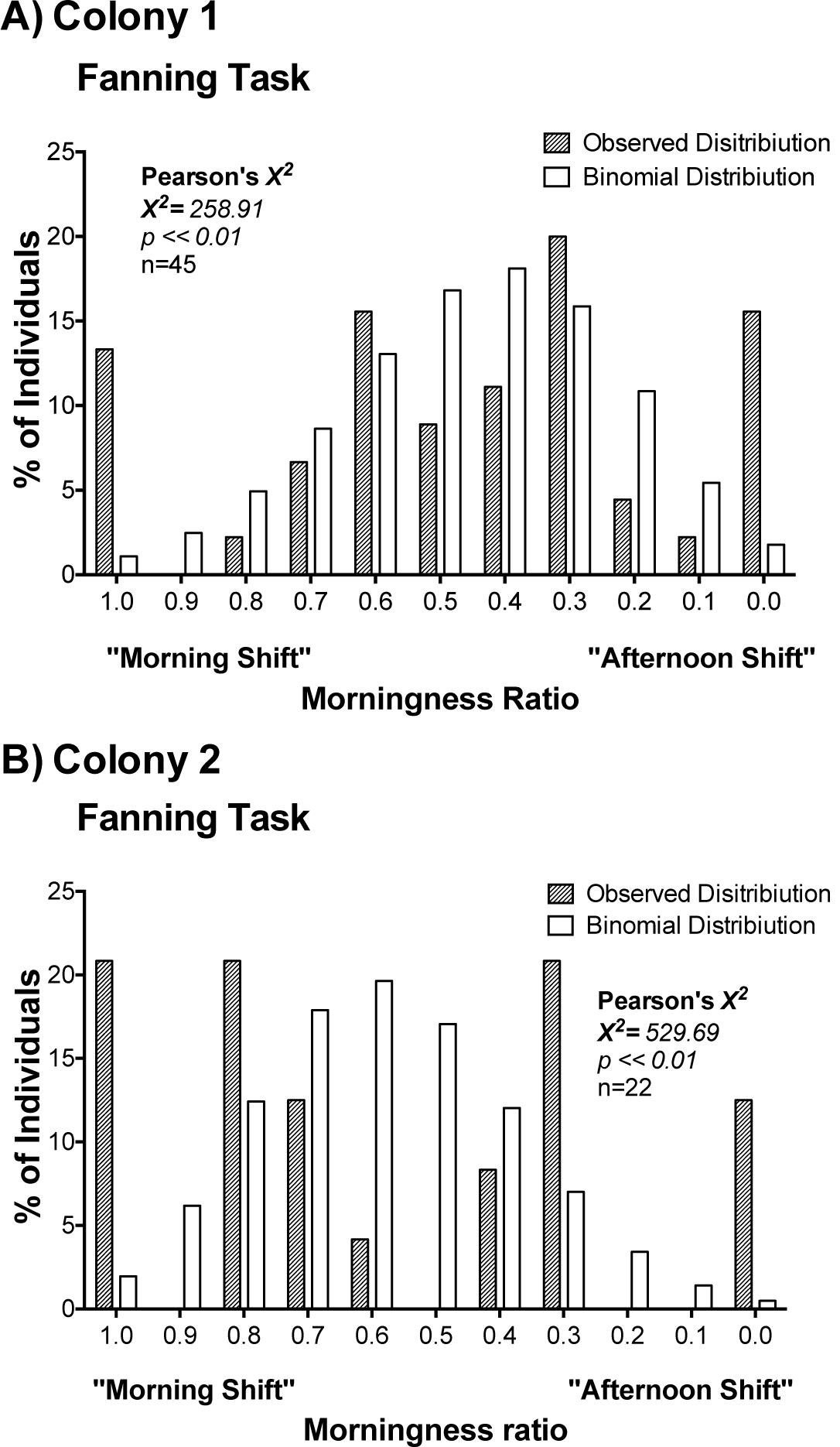
Exclusive morning and afternoon shifts are present in fanning task. **A)** Frequency distribution of the observed morningness ratio for colony 1 (shaded bars) of fanning behavior in the observation ramp. The observed distribution was compared to a theoretical binomial distribution (white bars) that assumes no shift work. Goodness of fit test revealed significant differences between the observed and theoretical distributions (*X*^*2*^*= 258*.*91, p<<0*.*01*, n=45). **B)** Frequency distributions of the observed morningness ratio (shaded bars) and theoretical binomial distribution (white bars for fanning behavior of colony 2. Consistent with the result from colony 1, Goodness of fit test showed significant differences between the observed and the binomial distribution (*X*^*2*^*=529*.*69, p<<0*.*01*, n=22). Comparison between the observed distributions for fanning behavior via Kolmogorov-Smirnov two-tailed test revealed significant differences between the frequency distributions of each colony (D= 0.346, *p<* 0.05).

### Endogenous relationship of foraging and fanning shifts

To examine how foraging and fanning shifts may be related we compared foraging and fanning morningness ratio of individuals that performed both foraging and fanning during our observations. This analysis resulted in a positive correlation between foraging and fanning shifts, suggesting that shift in one behavior influences the shift in the other (Figure 5A). However, upon closer inspection we observed that there were individuals had a shift for foraging but not for fanning and vice versa. Individuals that present shifts in both foraging and fanning behavior or lacked shifts were classified as presenting a non-dissociable shift, while individuals with shift in either foraging or fanning behavior were classified as dissociable shifts. By doing this we found that ∼30% of individuals present a dissociable shift (Figure 5B). These results suggest that foraging and fanning shifts are processes that are connected yet dissociable.

**Figure 5.**
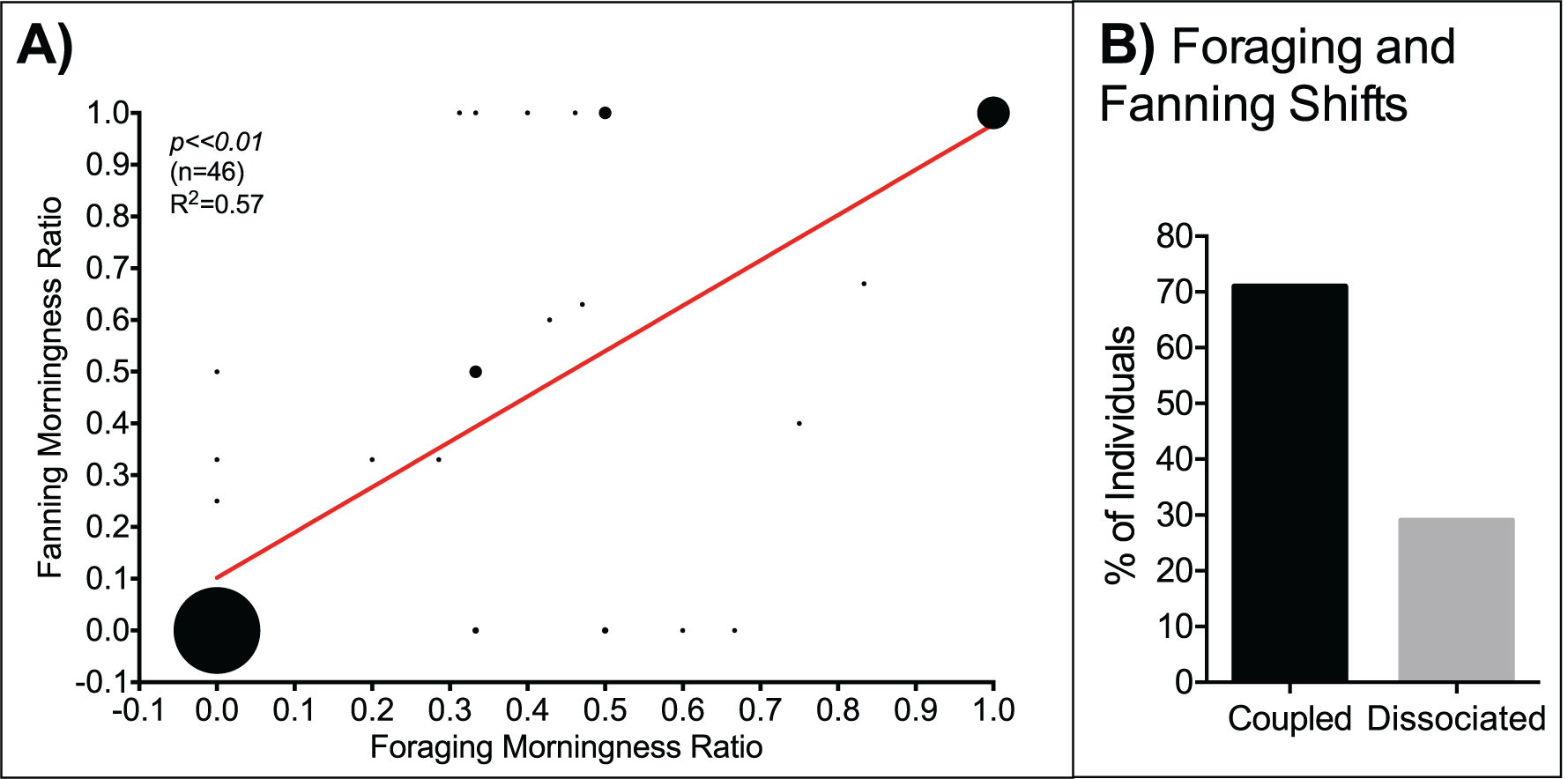
Foraging and fanning shifts are coupled yet dissociable behaviors. **A)** Pearson correlation of foraging and fanning morningness ratios for individuals that performed both tasks resulted in a positive correlation (R^2^=0.57, p<<0.01, n=46). The size of the dots is representative of the number of individuals in each data point. **B)** Per cent of individuals who’s foraging and fanning correlates (coupled) compared with those that do not correlate (dissociated).

## Discussion

The most significant finding of this study is that different shift work strategies contribute to the organization of different jobs in the honeybee colony. Foraging bees take advantage of the full daylight period to collect resources for the colony (Moore & Doherty, 2009; Moore & Rankin, 1983; Byron N. Van Nest & Moore, 2012; von Frisch, 1967; Wagner et al., 2013; Mark L Winston, 1987). Before our findings, it was not clear if foraging was performed continuously by each individual throughout the day or if distinct sub-groups (shifts) foraged at different times of the day. Here we show that both of these strategies are present in honey bee colonies, i.e. there are foragers that constantly work throughout the daylight period (Constant workers) and groups of foragers that only work in the morning or the afternoon (Shift workers) (Figure 1). In addition, we characterize various features of the honeybee shift work strategy. We observed that the demography of shift workers varies from task to task (Figure 2) and can be divided into individuals that maintain the same shift as they age, and those that change from one shift period to another (Figure 3). We also show that fanning, another task performed by workers, has a similar composition, with some individuals performing the job constantly throughout the day, and others doing so in shifts (Figure 4). Interestingly, around 60% of individuals, that were observed foraging and fanning, showed the same shift for both behaviors (Figure 5), suggesting that the shifts are coupled yet dissociable from one another.

We found that more than 40% of the individuals perform foraging trips exclusively in either the morning or the afternoon, while the remaining individuals (constant workers) forage throughout the daylight period (Figure 1). A previous study exploring the temporal organization of brood care found that nurses work around the clock (Moore et al., 1998). Their finding is consistent with the lack of circadian rhythmicity of nurses in the colony and the constant demand of brood care, regulated by brood pheromones (Yair Shemesh et al., 2010). In contrast, foragers are thought to rely on their circadian rhythms and time memory to successfully collect different resources and return to the colony (Moore & Doherty, 2009; Byron N. Van Nest & Moore, 2012; Wagner et al., 2013). The presence of both types of foragers (constant, shift workers) may be adaptive to the colony, and it could potentially result in the daylight period being more efficiently utilized by foragers.

Consistent with our hypothesis that the majority of foraging would be performed by constant workers, our results show that constant workers perform the majority (∼80%) of foraging trips (Figure 1). We expected that constant workers would perform at least twice the number of foraging trips than shift workers. By taking into account that we had 2 observation periods, that constant workers will be observed at both periods, and the proportion of shift workers and constant workers in our sample, we expected that constant workers would perform ∼75% of foraging trips. Since the predicted proportion of trips was similar to the predicted value (75% predicted vs. 80% obtained), the observed differences in workload between shift workers and constant workers can be accredited to 1) the higher proportion of constant workers and 2) the two potential observation periods for constant workers. It is likely that shift work was not uncovered directly until now since the majority of studies examining foragers at the colony or in artificial feeders make observations throughout the day, and until recently did not identify each individual. This combined with the low percentage of foraging flights taken by shift workers would significantly reduce the probability of collecting and observing shift workers in previous experimental setups.

Since honey bee foragers match their foraging activity to the time when the resource they are collecting is at the peak of production and establish a time memory of this event that allows them to anticipate resource availability (Moore & Doherty, 2009; Moore & Rankin, 1983; Moore, Siegfried, Wilson, & Rankin, 1989; Moore et al., 2011; Byron N. Van Nest & Moore, 2012; Wagner et al., 2013), it is possible that shift workers and constant workers visit groups of resources that are available at different times during the day. Evidence supporting this comes from the fact that, the temporal availability and duration of a resource, such as nectar or pollen, varies from flower to flower (Kleber, 1935; Linnaeus, 1755; Parker, 1926; von Buttel-Reepen, 1903). In addition, bees foraging to a food source that is available at noon or late in the afternoon have been shown to scout the food source on average up to 4 hours, prior to the resource availability on earlier days (Moore & Doherty, 2009; Moore & Rankin, 1993; Moore et al., 1989). Furthermore, once the resource a forager is exploiting closes for the day, the forager goes into the hive and does not take additional foraging flights for the day (Körner, 1940; Moore et al., 1989; Seeley, 1995; von Buttel-Reepen, 1903; von Frisch, 1940). It is possible that constant workers in our study are foraging to food sources available early in the afternoon, while afternoon shift workers are foraging to food source available in the late evening, but further studies are needed to test this hypothesis.

Alternatively, constant foragers could be classified as reticent foragers, who wait in the dance floor for a food source to be announced and forage as recruited by other individuals (Moore et al., 2011; Wagner et al., 2013). Another possibility is that the observed shift work strategy stems from the availability of stable food sources around the colony. In this scenario, foragers could specialize to more efficiently exploit a particular food source at its highest production point of the day, thus encouraging a shift work strategy. In contrast, a habitat where resources are scarce and constantly changing would foster foragers taking foraging trips at all times. Evidence for this notion stems from studies of different honey bee subspecies in Turkey, where *Apis mellifera syriaca*, which originate from an arid habitat with mild winters, presented higher flower fidelity than *A*.*m. carnica* and *A*.*m. caucasica*, which inhabit mountain regions with cold winters and short summers (Cakmak et al., 2010). Since the experiments in the previous study were performed in the same location it is likely that flower fidelity has a genetic component and this component may play a role in the shift work strategy that we observe in the current study.

Genotyping efforts by Kraus and colleagues (2011) suggested that shift work might be present and strongly affected by patrilineal genotype. Our findings are consistent with their measures, as pollen foragers with shift make up approximately 8% of the observed pollen specialists (Figure 2A). In contrast, approximately 36% of non-pollen foragers observed presented either a morning or afternoon shift (Figure 2B). This difference in the proportion of shift workers could be the result of intrinsic factors that differentiate pollen and non-pollen foragers, environmental factors such as resource availability or a combination of both. Studies examining resource specialization in foragers demonstrate intrinsic differences between pollen and nectar foragers, such as genetic background, sucrose responsiveness, phototaxis and octopamine titters (Barron et al., 2007; Erber et al., 2006; Giray et al., 2007; Page & Erber, 2002; Scheiner et al., 2003, 2001, 2002, 2014; Taylor et al., 1992; Wagener-Hulme et al., 1999). Given the similarity of proportion of shift work in this and the Kraus et. al study, it is likely that shift work in foraging may be dependent on foraging specialization. Alternatively, since pollen and nectar availability varies throughout the day from one flower to another (R.M. Goodwin, 1986; Linnaeus, 1755; Nakamura & Seeley, 2006; Stone, Willmer, & Alexandra Rowe, 1998), it is possible that the difference between pollen and non-pollen foragers stems from the availability of the particular resource a forager exploits. Future studies will examine how resource availability affects foraging timing and strategies and will explore if differences in patrilineal origin of non-pollen foragers influences their foraging shift.

We found that a group of individuals may begin foraging in either the morning or afternoon shift and over time switch shifts (Figure 3A). This switch is more probable to occur early in the foraging life (Figure 3C). This mechanism may be linked to epigenetic, hormonal, developmental or morphological changes occurring after the onset of foraging behavior (Brown, Napper, & Mercer, 2004; Farris, Robinson, & Fahrbach, 2001; Withers, Fahrbach, & Robinson, 1995). Since honeybee colonies need to constantly adapt to changes in the outside environment and resource availability, having a foraging force that can adjust at a moment’s notice may result in a constant flow of resources into the colony. Alternatively, it is possible that changes in the timing of foraging result from the disappearance of the resource the bee was exploiting, causing her to visit a new resource that may be available at a different time. Although much work remains to be done, both of these scenarios are consistent with the idea that shift work may be plastic and thus adopts to the colony’s constant needs.

While our findings show that some individuals perform foraging in shifts, our direct observations of foraging behavior cannot determine if shifts are intrinsic or a function of external factors. While assaying foraging we also observed fanning at the entrance of the colony. To our surprise, we found that some individuals fanned exclusively in the morning or afternoon, while others had no preference for a specific shift (Figure 4). The observed shifts in fanning suggest that shift work may have one or more intrinsic drivers. One of these drivers may be genetic variation among individuals in the colony, as previously described for pollen foragers (Kraus et al., 2011). Previous studies looking at genetic variation within fanning bees found that colonies with natural genetic variation have a more rigorous control of temperature inside of the colony (via fanning) in comparison with colonies that originate from a single artificially inseminated queen (J C Jones et al., 2004; Julia C Jones, Nanork, & Oldroyd, 2007; Su et al., 2007). Furthermore, evidence suggests that genetic variation in the colony increases overall colony fitness (Mattila & Seeley, 2007). Taken together, our data on fanning task and that of previous studies, it is possible that shift work in honeybees has one or more intrinsic mechanisms driving it. If this driver or drivers have a genetic component, the study of single-cohort colonies may result in the loss of one or both shifts in foraging and fanning tasks.

Since some of the marked individuals we observed foraging also fanned, we explored the potential relationship of shift work between these tasks. Our results revealed that while a proportion of individuals (30%) perform foraging and fanning behaviors at different time periods, the remaining individuals presented the same shift for both foraging and fanning (Figure 5). This suggests that while these tasks may share a relationship with regards to shifts, they can be dissociated from one another (Figure 5B). This difference between foraging and fanning shifts could be explained by differences in the influences of endogenous (genetic background, life stage) and exogenous factors (light, temperature, resource availability, colony needs). Previous work done using the fruit fly *Drosophila melanogaster*, uncovered experimental proof of the multiple circadian oscillator hypothesis originally proposed by Dan and Pittendrigh (Colin S. Pittendrigh & Daan, 1976; Stoleru, Peng, Agosto-Rivera, & Rosbash, 2004). This hypothesis states that complex multicellular organisms possess various independent or loosely coupled circadian pacemakers (C. S. Pittendrigh, 1972). In the case of the fruit fly researchers uncovered that different cells were responsible for the morning and evening activity peaks in locomotor behavior (Stoleru et al., 2004). Similarly, we hypothesize that each task (foraging and fanning) is under a set of different circadian oscillators and while the oscillators may be in synchrony in some individuals, this may vary across individuals.

Based on our observation of a shift work strategy in foraging and fanning tasks, we posit that the use of this strategy may confer various benefits to honeybee colonies. The use of shift work in foraging will allow the colony to take advantage of stable resources available throughout the day. Constant workers could enhance the efficiency of shift workers by being ready to forage when a food source is announced, thus increasing the number of foraging flights to the particular food source. Although having shift work can provide a number of benefits to the hive, it may have negative effects on the individual. For example, if we presume that phase differences in the circadian clock underlie the observed shift work, then some individuals may be desynchronized with respect to environmental cycles. A number of studies in humans have shown that individuals with evening chronotypes have increased susceptibility to a number of disorders such as circadian misalignment, cancer and depression (Adan et al., 2012; Antunes, Levandovski, Dantas, Caumo, & Hidalgo, 2010; Davis & Mirick, 2006; Dibner, Schibler, & Albrecht, 2010; Lépine & Briley, 2011; Reinberg, Touitou, Lewy, & Mechkouri, 2010). In the case of honeybees, shift work could potentially have negative effects on the individual workers. Future studies will look at dissecting the relationship between shift work in foraging and fanning behavior and circadian rhythms in bees (Giannoni-Guzmán et al., 2014).

In conclusion, this study shows for the first time direct behavioral evidence of shift work strategy being used in foraging and fanning tasks in honeybee colonies and characterize behavioral components of this shift work strategy. These findings reveal yet a new layer of social and temporal organization of honeybee colonies. Future studies may aim to understand the specific genetic components and neural mechanisms underlying shift work. Since honeybees use their endogenous circadian clock to predict time of day, the relationship between circadian rhythms and shift work is an area of great interest. Studying this relationship may eventually provide clues on how to attack the negative consequences of imposed shift work in humans.

## Acknowledgements

We would like to thank honeybee specialist Gabriel Diaz for honeybee colony care. Dr. Arian Avalos, Emmanuel Rivera, Liz Hernandez, Jonathan Aleman-Rios and Carlos Vazquez for help with the experiments. Thanks to Dr. Arian Avalos and Luis de Jesus for their comments and suggestions. We would also like to recognize the director, Manuel Diaz and the personnel of the Gurabo Experimental Agriculture Station of the University of Puerto Rico at Mayaguez for use of facilities at ‘‘Casa Amarilla’’. This work was sponsored by the National Science Foundation (NSF) awards 1026560, 1633184, 1707355 and the National Institute of Health (NIH) 2R25GM061151-13, P20GM103475.

